# Intrinsic Functional Connectivity in the Default Mode Network Predicts Mnemonic Discrimination: A Connectome-based Modeling Approach

**DOI:** 10.1101/2021.07.09.451777

**Authors:** Christopher N. Wahlheim, Alexander P. Christensen, Zachariah M. Reagh, Brittany S. Cassidy

## Abstract

The ability to distinguish existing memories from similar perceptual experiences is a core feature of episodic memory. This ability is often examined using the Mnemonic Similarity Task in which people discriminate memories of studied objects from perceptually similar lures. Studies of the neural basis of such mnemonic discrimination have mostly focused on hippocampal function and connectivity. However, default mode network (DMN) connectivity may also support such discrimination, given that the DMN includes the hippocampus, and its connectivity supports many aspects of episodic memory. Here, we used connectome-based predictive modeling to identify associations between intrinsic DMN connectivity and mnemonic discrimination. We leveraged a wide range of abilities across healthy younger and older adults to facilitate this predictive approach. Resting-state functional connectivity in the DMN predicted mnemonic discrimination outside the MRI scanner, especially among prefrontal and temporal regions and including several hippocampal regions. This predictive relationship was stronger for younger than older adults, primarily for temporal-prefrontal connectivity. The novel associations established here are consistent with mounting evidence that broader cortical networks including the hippocampus support mnemonic discrimination. They also suggest that age-related network disruptions undermine the extent that the DMN supports this ability. This study provides the first indication of how intrinsic functional properties of the DMN support mnemonic discrimination.

## Introduction

Encoding and retrieving everyday events allows people to relive past experiences and plan future behaviors. Because people experience countless overlapping events across their lifetimes, they must be able to distinguish existing memories from sensory inputs. For example, someone could leave the table during dinner and return to find a similar glass that is not theirs. One way they could avoid drinking from the similar glass is to discriminate it from their memory for the original glass. The subsequent interference created by such feature similarities can be mitigated when one encodes the inputs (i.e., the glasses) as distinct representations. Such encoding would allow one to avoid future confusion about the ownership of both glasses because the unique features of each would be represented with distinguishing associative information. This example of interference reduction may be accomplished by a hippocampal computation referred to as pattern separation that is enabled by the orthogonalization of overlapping inputs (1–3). Evidence for pattern separation computations has been inferred from studies showing that hippocampal function and structure predicts mnemonic discrimination (for a review, see 4).

Substantial evidence for this relationship has been observed using modified recognition memory paradigms, such as the Mnemonic Similarity Task (MST; for a review, see (4)). In the object variant of the MST, participants study everyday objects (e.g., a glass) and complete a memory test comprising exact repetitions of studied objects (e.g., the same glass) and lures that are similar but not identical to studied objects (e.g., a shorter glass). Mnemonic discrimination occurs when lures are identified as similar but not identical to studied objects. Functional magnetic resonance imaging (fMRI) studies show that such discrimination reliably evokes activity in hippocampal subfields (e.g., dentate gyrus and CA3) in paradigms in which the similarity among materials during study and test is manipulated for objects and scenes (5–8), spatial information (9, 10), and temporal intervals (9).

Although most studies of the neural correlates of mnemonic discrimination have focused on the role of hippocampal subfields, there is mounting evidence that cortical regions are also involved (10–13). Recent fMRI studies have begun to characterize activation outside the hippocampus that supports mnemonic discrimination by examining whole-brain activity during the MST. For example, functional connectivity from the hippocampus to bilateral temporal regions as well as the cerebellum and frontal and temporal cortex has been shown during mnemonic discrimination (14). Other work has shown that activation in hippocampus as well as prefrontal and occipital cortex was associated with mnemonic discrimination (15), with the latter suggesting contributions of visual information that are not directly involved in discrimination (also see, 16). Further, brain activity during mnemonic discrimination has been shown in regions of the default mode network (DMN; 17), including the precuneus and angular gyrus (18), that is coupled with hippocampal activation during mnemonic discrimination. Finally, inactivation of anterior DMN regions (e.g., medial prefrontal cortex) in rodents impairs mnemonic discrimination (19). These studies suggest that regional DMN connectivity may support mnemonic discrimination.

This suggestion is also supported by studies showing that episodic memory performance is associated with task-related activity in DMN regions. Such activation has been observed in regions including the precuneus, angular gyrus, temporopolar cortex, and medial prefrontal cortex (20, 21). These regions of the DMN have also been shown to operate with the hippocampus, a core structure involved in mnemonic discrimination, during episodic memory retrieval (22). Although there is not a universal consensus that the hippocampus is formally part of the DMN (17), it is densely interconnected with many DMN regions that support episodic memory (23).

A role for DMN connectivity in mnemonic discrimination ability may be identified by examining how intrinsic functional connectivity at rest predicts task performance. Such brain-behavior relationships are suitable for this purpose because they identify which networks support specific cognitive functions (24) and are consistent with the functional network architecture revealed during tasks (25). This consistency between resting-state and task-related approaches inspired the view that task-evoked activity over intrinsic brain networks is a mechanism that supports cognitive operations (e.g., 26). Accordingly, intrinsic DMN connectivity associated with episodic memory (27, 28) may support mnemonic discrimination. This may occur when such functional neural architecture enables a recall-to-reject mechanism (1) to access stored representations and compare them with sensory inputs. From this view, the connectivity among episodic memory regions should mediate mnemonic discrimination to the extent that they support access to high-fidelity memory representations.

If intrinsic DMN connectivity supports mnemonic discrimination, then this relationship should also be observed across a wide range of performance. One way to test this assertion is to examine whether this association occurs across people from groups that consistently differ in mnemonic discrimination, such as healthy younger and older adults (e.g., 29. Testing people across age groups thus enables examination of a key source of variability in mnemonic discrimination ability. Age-related mnemonic discrimination deficits have been linked to hyperactivity in the hippocampus (3, 30– 32), hypoactivity in MTL cortical regions (30, 31), and dysfunctional connectivity between anterior hippocampus and parahippocampal cortex (33). These findings suggest that network-level perturbations in aging may extend beyond the MTL into networks supporting memory, such as the DMN. An ideal method for examining the association between intrinsic functional connectivity of the DMN and mnemonic discrimination is Connectome-based Predictive Modeling (CPM; 34). CPM is a data-driven, cross-validation approach that builds a connectome associated with a specific behavior (e.g., mnemonic discrimination). A connectome is a set of functionally connected regions of interest (ROIs) used to predict the behavior of novel participants. CPM is favored over many related approaches, partly because it effectively predicts behavior using the connectivity between all ROIs without hypotheses about which connections will contribute (34). CPM has been used to identify whole-brain connectivity patterns at rest and during tasks that measure fluid intelligence (35), attentional control (36, 37), creative ability (38), and personality (39). It has also been used to characterize aspects of older adults’ cognition (40). Together, this nascent literature suggests that the CPM approach is well-suited to explore the extent that intrinsic functional connectivity in DMN regions predicts a wide range of mnemonic discrimination abilities.

## The Present Study

Our overarching goal was to characterize associations between intrinsic functional DMN connectivity and mnemonic discrimination for the first time. The recent introduction of CPM affords a unique opportunity to characterize such associations using a method that could eventually generalize to predict behaviors in other data sets. Participants first provided resting-state fMRI data and, outside the scanner, completed an object-based MST (29) followed by a perceptual discrimination task (PDT). Including both tasks enabled the isolation of relationships between neural activation and mnemonic processes in the following aims.

Our primary aim was to provide the first characterization of the association between intrinsic DMN connectivity and mnemonic discrimination using the CPM approach. Consistent with studies showing that the functional architecture of brain networks in comparable in resting and task states (25, 26), previous work using CPM has demonstrated that resting state connectomes are associated with task-based connectomes (36). Given the findings above indicating that intrinsic DMN connectivity is associated with episodic memory, we expected that DMN connectivity would predict mnemonic discrimination. Our rationale is that mnemonic discrimination can be accomplished when similar lures trigger episodic retrievals of and comparisons with representations of studied objects. As described above, this processing sequence can unfold when participants deploy a recallto-reject strategy meditated by hippocampal operations (1).

To further connect with studies examining hippocampal contributions to mnemonic discrimination in the MST (for a review 4), we included in the CPM distinct regions for the hippocampal head, body, and tail. Our exploratory aim here was to determine if predictive inter-region connections differed along the hippocampal longitudinal axis. Our approach was inspired by findings showing that anterior and posterior regions are differentially associated with encoding and retrieval processes (41) as well as memory for gist and item-specific information (42), respectively. We were also inspired by the view that cortical regions supporting memory for items and their context are differentially connected with anterior and posterior hippocampal regions (for a review, see 23). Additionally, some evidence suggests that anterior and posterior hippocampus may be differentially vulnerable in aging, though there are somewhat conflicting results about whether anterior (33) or posterior (43, 44) hippocampal regions show more pronounced age-related change. Despite these differential associations across hippocampal regions, the absence of prior work applying CPM to episodic memory abilities preclude theoretically motivated a priori hypotheses about how connectivity profiles may differ across those regions. Therefore, we report the first characterization of such profiles below and consider the implications of such connectivity further in the Discussion.

Our secondary aim was to examine whether the extent to which DMN connectivity predicts mnemonic discrimination differs between younger and older adults. Older adults consistently show impaired mnemonic discrimination (for a review, see 4) and weakened intrinsic DMN connectivity associated with episodic memory deficits (45, 46). Consequently, weaker intrinsic DMN connectivity in older adults may undermine the extent that DMN connections predict their mnemonic discrimination ability in the CPM. We therefore hypothesized that intrinsic DMN connectivity in the CPM would predict mnemonic discrimination more strongly for younger than older adults.

## Method

The materials, anonymized data files, coded recall responses, and analysis scripts are available on the Open Science Framework (OSF): https://osf.io/f6vg8/. The study was approved by the University of North Carolina at Greensboro (UNCG) Institutional Review Board.

### Participants

The participants were 36 younger and 36 older adults from the greater Greensboro, North Carolina community. They were right-handed and had no recent history of neurological problems. The stopping rule was to collect usable data from at least 25 participants per age group. This sample size is comparable to research examining associations between functional connectivity and mnemonic discrimination in younger and older adults (33). We excluded one younger and one older adult who scored below 26 on the Montreal Cognitive Assessment (MoCA; 47). We also excluded one younger adult for excessive movement in the scanner, five older adults for responding on fewer than 70% of the MST trials, and two older adults for not following instructions. The final sample included 34 younger adults (18-32 years old, *M*_*age*_ = 22.21, *SD* = 3.65; 20 female) and 28 older adults (61-80 years old, *M*_*age*_ = 69.82, *SD* = 5.64; 20 female). Table 1 shows that younger and older adults had comparable education and working memory capacity, the latter measured by forward and backward digit span (48). Older adults had higher vocabulary scores (49) and slower processing speed (50) than younger adults.

**Table 1.** Demographic Information and Cognitive Ability Scores for Younger and Older Adults

|  | Younger | Older | *t* | *p* | Cohen's *d* |
| --- | --- | --- | --- | --- | --- |
| Years of education | 15.85 (2.50) | 15.75 (2.29) | 0.17 | = .867 | 0.04 |
| Vocabulary | 29.03 (3.97) | 33.64 (3.12) | 5.01 | < .001 | 1.29 |
| Processing speed | 78.26 (13.27) | 63.36 (3.12) | 4.51 | < .001 | 1.15 |
| Digit span (forward) | 8.71 (2.14) | 8.00 (2.09) | 1.31 | = .196 | 0.33 |
| Digit span (backward) | 7.38 (2.22) | 6.82 (1.74) | 1.09 | = .280 | 0.28 |
| MoCA | 28.26 (1.42) | 27.68 (1.39) | 1.63 | = .108 | 0.41 |

*Note.* Means and standard deviations (in parentheses) are displayed above. Vocabulary was quantified via the Shipley Vocabulary Test; Processing Speed was quantified via Digit Comparison Task; digit span scores were measured via tasks included in the Wechsler Memory Scales.

### Behavioral Tasks

Participants were tested individually in a quiet room. An experimenter explained the instructions before each task and then allowed participants to complete the task alone. Stimuli were presented electronically using EPrime 3.0 software (51) on a PC laptop that included a 13.3 in (33.78 cm) display (1920 × 1080 resolution). The viewing distance was approximately 20 in (51 cm). Images of everyday objects (400 × 400 pixels) were from a publicly available database (https://github.com/celstark/MST).

### Mnemonic Similarity Task

Participants first completed a version of the object-based MST (Figure 1A) that included separate study and test phases. During study, pictures of everyday objects appeared in the center of the display against a white background for 2000 ms each. Participants pressed a key to indicate if the object belonged indoors (v) or outdoors (n). At test, participants viewed three object types for 2000 ms each. Test objects repeated studied objects exactly (repeated objects), were similar but not identical to studied objects (similar lures), or had not appeared during study (novel foils). Participants pressed a key to classify repeated objects as “old” (v), similar lures as “similar” (b), and novel foils as “new” (n). Objects appeared in random order in each phase. Each participant viewed 72 study objects and 108 test objects. The test included 36 *repeated objects*, 36 *similar lures*, and 36 *novel foils*. These object type conditions included fewer items than is typically used in the MST; however, performance on key measures has been shown to be consistent with even fewer items (i.e., 16 per condition; 29). For all object types, each 36-object set included 12 objects from each of the three “lure bin” sets that created the greatest challenge for mnemonic discrimination by including the most confusable lures (i.e., bins 1-3). Lure bins are pre-established object groupings based on probabilities of misclassifying similar lures as repeated objects, with lower-numbered bins indicating more of such false alarm errors (29). These error rates were equated across sets. For counterbalancing, sets were rotated through conditions, creating three experimental formats.

**Fig. 1.**
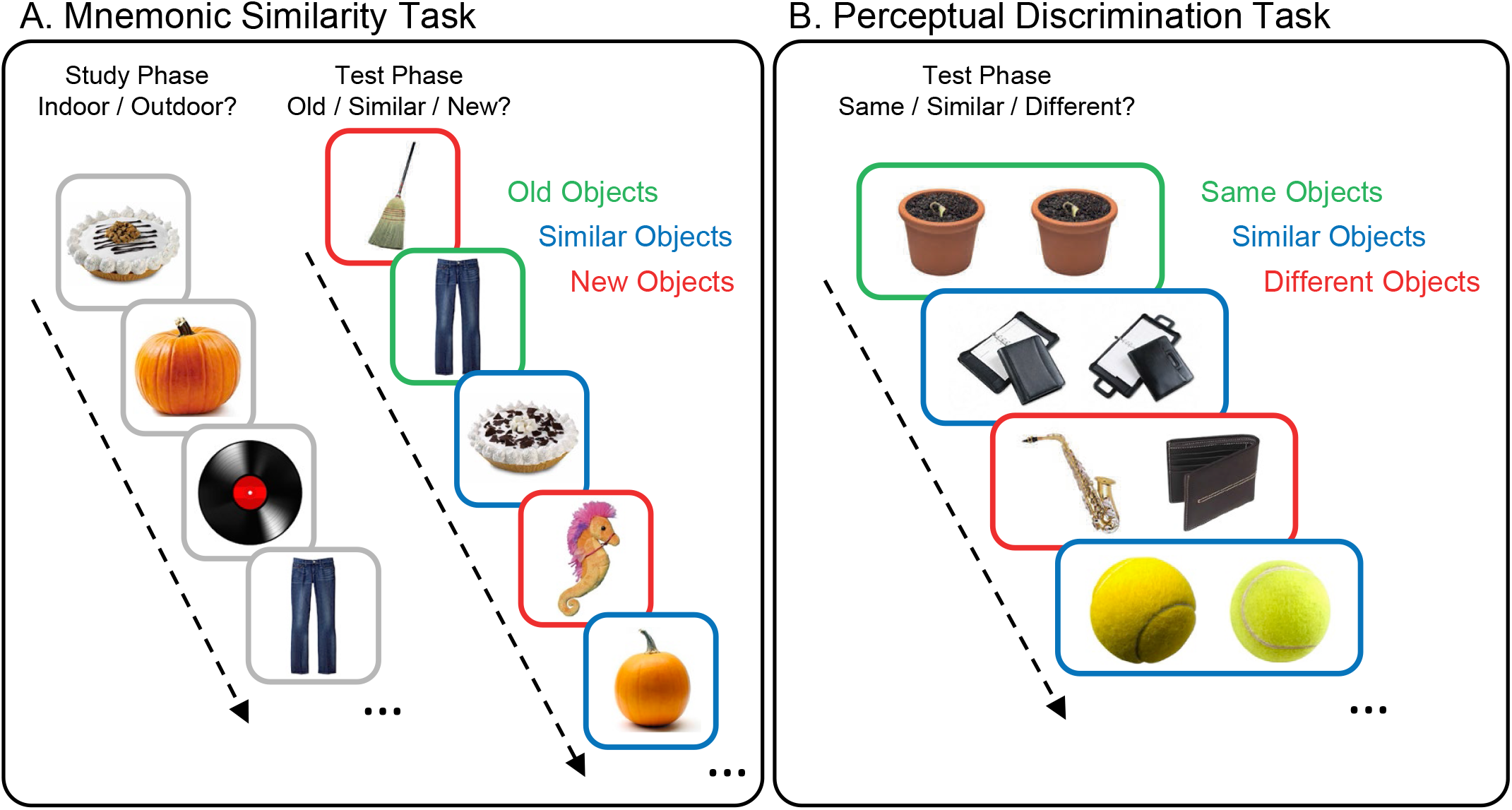
Schematics of Behavioral Tasks. (A) In the Mnemonic Similarity Task, participants studied pictures of objects and took a recognition test in separate phases. The recognition test included old objects that were repeated from the study phase, similar objects that were alternative versions of other old objects from the study phase, and new objects that only appeared in the test phase. During study, participants indicated whether each object belonged indoors or outdoors. At test, participants indicated whether objects were old, similar, or new. (B) In the Perceptual Discrimination Task, two objects appeared side-by-side on each trial. Some pairs included two pictures of the exact same object, others included two similar versions of the same object, and another set included two different objects. Participants responded to each pair by indicating whether they included the same, similar, or different objects.

Note that the combination of shorter lists and more confusable lures relative to some variants of the MST using object stimuli could have offsetting effects on levels of mnemonic discrimination scores. The shorter study list here compared to earlier studies could have led to better recognition of studied objects via a list-length effect occurring when shorter lists improve memory (for a review, see (52)). Such improved recognition could also have supported a recall-to-reject strategy of similar lure identification. However, list-length effects are not universal and are absent when stimulus features support inter-item discriminability (e.g., 53. Also, including only the most confusable lures here should challenge the comparison process of the recall-to-reject strategy, thus reducing mnemonic discrimination. These offsetting influences suggest that the observed mnemonic discrimination scores reported below could be higher than, comparable to, or lower than scores in other studies. But the exact balance of these influences can only be determined via an experimental comparison that we did not conduct here. Regardless of how the present paradigm compares to others, it effectively served as a time-efficient task with an ideal range of performance for CPM analyses (see below).

### Perceptual Discrimination Task

After the MST, participants completed the PDT (Figure 1B). The PDT was included to account for perceptual ability in mnemonic discrimination differences (see 54). The PDT included different objects than the MST to prevent cross-task contamination. Normative false alarm error rates for similar lure objects were equated across the tasks to control for stimulus effects. Participants were instructed to classify relationships within pairs. Two objects appeared together on each trial until participants responded. *Repeated Objects* included identical objects (e.g., same potted plant); *Similar Objects* included two similar but not identical versions of the same object (e.g., similar planners); and *Novel Objects* included objects with different identities (e.g., saxophone and wallet). Participants pressed a key to classify pairs as comprising the “same” objects (v), “similar” objects (b), or “different” objects (n), for repeated objects, similar lures, and novel objects, respectively. Pairs appeared in random order. There were 90 total pairs (36 repeated objects, 36 *similar lures*, and 18 *novel objects*) with 18 unique object identities per condition (54 identities). For counterbalancing, object sets were rotated through conditions, creating three experimental formats.

### Statistical Approach for Assessing Behavioral Task Performance

The analytic approach for the MST followed prior studies in that response probabilities for each object type and bias-corrected mnemonic discrimination and traditional recognition indices are reported (4). The mnemonic discrimination index, referred to as the Lure Discrimination Index (LDI), was calculated as the difference in “similar” responses to lures and foils: p|similar (lures – foils). In addition, the traditional recognition index was calculated as the difference in “old” responses to repetitions and foils: p|old (repetitions – foils). The analytic approach for the PDT is comparable in that response probabilities and a bias-corrected index of perceptual discrimination are reported. A perceptual discrimination index was calculated as the difference in *similar* responses between similar object pairs and novel object pairs: p|similar (pairs with similar objects – pairs with novel objects). All analyses of behavioral performance were conducted using R software (55). Pairwise comparisons of the bias-corrected measures for younger and older adults were conducted using the emmeans function from the *emmeans* package (56). Analysis scripts are available on the OSF: https://osf.io/f6vg8/. The level for significance was set at *α* = .05.

### fMRI Data Acquisition and Preprocessing

Before completing the behavioral tasks, but after completing the neuropsychological measures, whole-brain imaging was performed on a Siemens 3.0T Tim Trio MRI Scanner using a 16-channel head coil at the Gateway MRI Center at UNCG. High-resolution anatomical images were acquired with a multi-planar rapidly acquired gradient echo (MP-RAGE) sequence (192 sagittal slices, 1 mm thickness, TR = 2300 ms, TE = 2.26 ms, 1.0 mm isotropic voxels). Restingstate functional data were collected after the anatomical scan. Resting-state data comprised 300 measurements collected over one 10-minute run. This scan duration produces reliable functional connectivity estimates within and across participants (57). Participants were instructed to remain still and awake with their eyes open. No stimuli appeared during this scan, and the display was black. Functional scans were collected using an echo-planar image sequence sensitive to blood-oxygen-level-dependent (BOLD) contrast (T2*; 32 slices with 4.0 mm thickness and no skip, TE = 30 ms, TR = 2000 ms, flip angle = 70, FOV = 220 mm, matrix size = 74 × 74 × 32 voxels, A/P phase encoding direction). Slices were collected in a descending order that covered the entire cortex and partial cerebellum. At the beginning of the resting-state scan, the scanner acquired and discarded two dummy scans. These structural and resting-state fMRI data are available on OpenNeuro: https://openneuro.org/datasets/ds003871/versions/1.0.2.

Data were preprocessed with the default preprocessing pipeline in the CONN functional connectivity toolbox (58) used in conjunction with SPM12 (Wellcome Trust Centre for Neuroimaging, London, UK; www.fil.ion.ucl.ac.uk/spm). This pipeline is described here (https://web.conn-toolbox.org/fmri-methods/preprocessing-pipeline) and is summarized in what follows. Within this pipeline, images are 1) realigned to correct for motion, 2) slice-time corrected, and 3) undergo outlier identification using custom artifact detection software (http://www.nitrc.org/projects/artifact_*d*_*etect*) that detects outlier time points for each participant. Outlier scans are identified from the observed global BOLD signal and participant motion in the scanner. Following recommended conservative parameters for examining associations between DMN connectivity and age differences in behavior (59), volumes were excluded if the signal for that time point fell three standard deviations outside the mean global signal for the entire run or if the scan-to-scan head motion exceeded .5 mm in any direction. Note that this software takes both scan-to-scan head motions and rotations into account when estimating composite motion (similar to framewise displacement).

Following outlier identification, images were normalized to the MNI template, resampled to 3 mm isotropic voxels, and smoothed using an 8 mm FWHM isotropic Gaussian kernel. This smoothing precludes biases associated with spatial registration to a template across age groups (e.g., 60). While smoothing to 8 mm may not be ideal for hippocampal long-axis dissociations, this is not an unusually large kernel (cf. 61–63. Moreover, consistency in template coregistration across the entire sample was a greater priority for these analyses than minimizing smoothing kernel size.

The preprocessed data were then subjected to the default denoising pipeline within CONN described here: https://web.conn-toolbox.org/fmri-methods/denoising-pipeline. This pipeline consists of a linear regression of potential confounding effects in the BOLD signal and temporal band-pass filtering. Potential confounding effects include noise components from cerebral white matter and cerebrospinal areas (64), estimated motion parameters identified during the realignment step of preprocessing, outlier scans from outlier identification (i.e., scrubbing; 65), and constant and first-order linear session effects (58). This approach preserves valid positive functional connectivity estimates while controlling for the inflation of negative estimates and regresses out physiological noise from areas of non-interest (e.g., white matter; see 64). The residual time series was then band-pass filtered in the 0.008–0.08 Hz range.

Stable correlations can be computed from resting-state functional data using around five minutes of cleaned data (65, 66). One younger adult was excluded for not meeting this threshold. The analyzed sample (34 younger and 28 older adults) showed patterns typical in aging and connectivity work (e.g., 59). There were more outliers scans for older adults (*M* = 34.71, *SD* = 29.15) than younger adults (*M* = 19.47, *SD* = 20.31), *t*(60) = 2.42, *p* = 0.019, *d* = 0.61. After removing these scans, younger adults retained an average of 9.36 / 10 minutes (*SD* = 0.67) and older adults retained an average of 8.84 / 10 minutes (*SD* = 0.97) of data. The final sample included enough data to calculate stable resting-state correlations (65, 66), thus mitigating concerns about analysis results being skewed by age differences in outlier scans.

Functional connectivity matrices were calculated across the time series using Fisher’s *z* coefficients between 200 cortical ROIs isolated using the Schaefer parcellation based on the MNI template (67) and eight hippocampal regions (left and right medial and lateral head, left and right body, and left and right tail) extracted from the Melbourne Subcortex Atlas using the 2 mm group parcellation (68). Because there was no a priori reason to include medial and lateral hippocampal head ROIs, the medial and lateral hippocampus Fisher’s *z* coefficients were averaged for left and right hemisphere. All *z* coefficient values were then transformed to Pearson’s *r* correlation coefficients for ease of interpretation. Functional connectivity matrices were then constrained to the 37 ROIs labeled in the parcellation as being part of the DMN and the six hippocampal regions—bilateral head, body, and tail—for the CPM analyses described below (Table 2 contains MNI coordinates for each ROI). There was thus one 43 × 43 functional connectivity matrix per participant. Note that there would be no difference in connectivity values when generating a connectivity matrix across all ROIs and then restricting an analysis to a subset relative to only generating a connectivity matrix restricted to a subset of ROIs. We chose the former to suit lab interests unrelated to the current research questions. These functional connectivity matrices are available on the OSF: https://osf.io/f6vg8/.

**Table 2.**
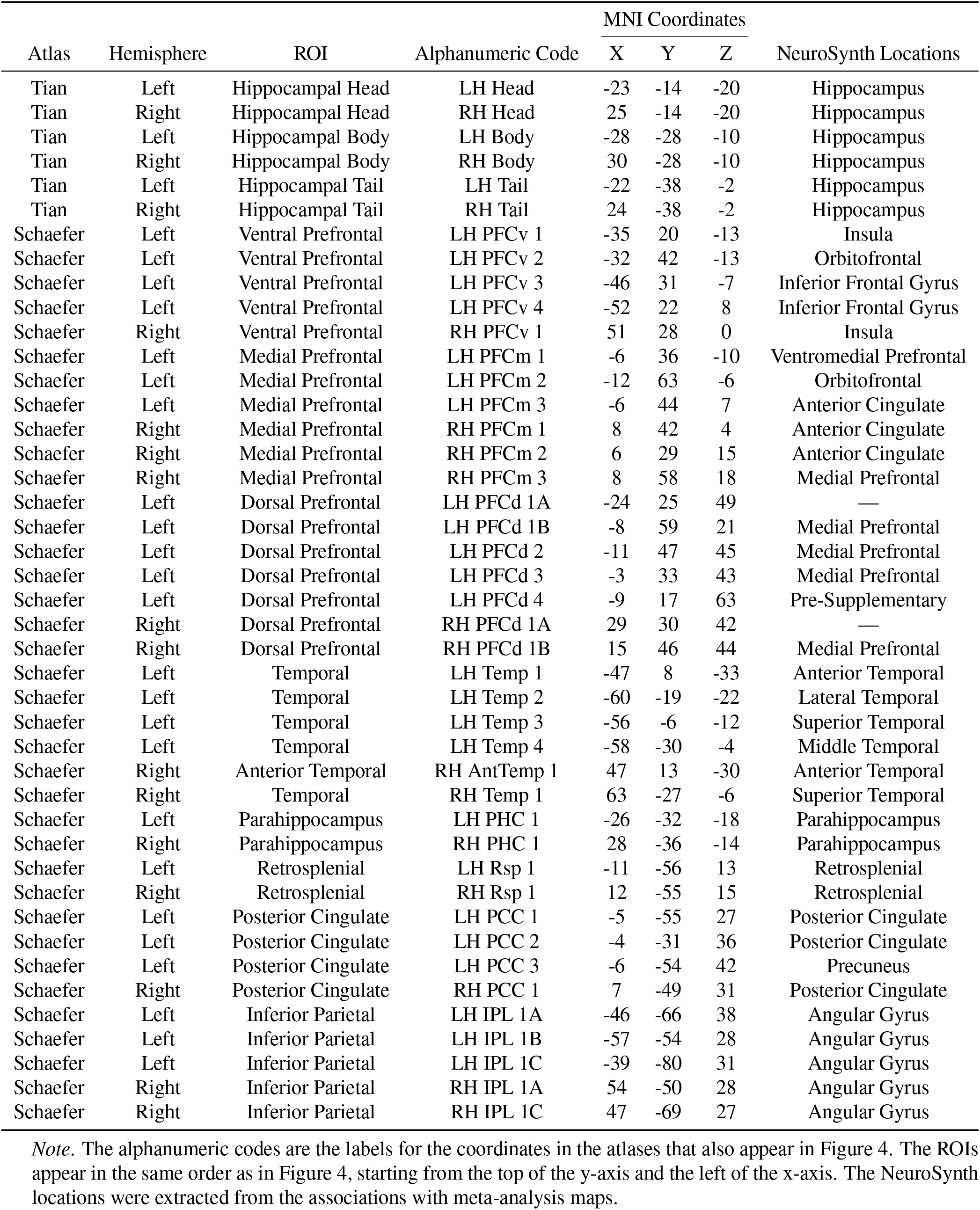
MNI Coordinates of Tian and Schaefer Atlases

### Statistical Approach for Generating Connectivity Matrices

CPMs were estimated using the *NetworkToolbox* package (version 1.4.2; 69) in R software. CPM constructs a connectome that is significantly associated with a behavioral measure via a leave-one-subject-out cross-validation approach (e.g., 35). Each behavioral measure used in CPM generates a specific connectome. Here, we generated a connectome for mnemonic discrimination using the LDI score as the behavioral measure. Each participant’s resting-state data comprised the functional connectivity matrix of ROIs described above, henceforth referred to as ROI–ROI matrices. The CPM approach proceeded as follows. First, the ROI–ROI matrix and mnemonic discrimination for one participant (i.e., the test participant) was separated from the remaining participants (i.e., training participants). The training participants’ ROI–ROI matrices were then used to correlate each ROI–ROI element across the matrices with their respective mnemonic discrimination performance. To rule out artifactual relationships from age-related structural brain differences, total intracranial volume, including gray matter, white matter, and cerebrospinal fluid, was controlled when computing correlations between ROI–ROI elements and LDI scores. These correlations formed a single correlation matrix thresholded at *α* = .05 (34).

Next, the thresholded matrix was separated into two masks that comprised only significant positive or negative correlations. Creating these separate masks allowed us to dissociate connectivity patterns that were positively and negatively associated with mnemonic discrimination. Each mask was separately applied to each training participant’s ROI–ROI matrix, leaving only the connectivity values that were significantly related to behavioral performance. For each participant, sum totals of these remaining connectivity values were calculated for each mask. These totals were then entered into separate regression models to predict the training participants’ mnemonic discrimination performance.

The regression weights from these models were then used to predict the test participant’s mnemonic discrimination. Using the test participant’s ROI–ROI matrix, sum scores from the positive and negative masks were computed to estimate mnemonic discrimination for the test participant by solving the respective regression equations. The predicted LDI value from the positive mask is referred to as the positive prediction, and the predicted LDI value from the negative mask is referred to as the negative prediction. This process repeated until all participants served once as the test participant, thus producing two predicted LDI values per participant. Finally, the connectome-predicted and observed LDI values were correlated. A significant correlation indicates that a set of connected ROIs (i.e., connectome) had sufficient explanatory power for mnemonic discrimination performance.

Note that our approach to constructing the ROI-ROI matrix used to establish a predictive model of mnemonic discrimination ability did not account for age group differences in intrinsic functional connectivity in the model. This approach was intentional because it allowed us to identify age-related differences in the extent to which particular ROI-ROI connections predicted mnemonic discrimination performance (see results below). But this approach could also lead to a connectome driven mostly by age differences instead of individual variation in mnemonic discrimination across the entire sample. To account for age differences in the predictive validity of the connectome, we conducted parallel analyses correlating the ROI-ROI matrix with behavioral performance but also included age as a continuous covariate (34, 39, see Supplemental Information, Section 1 [SI1]).

## Results

### Behavioral Tasks

The MST response probabilities are displayed in SI Table S1 (top rows). Mnemonic discrimination (Figure 2A) was significantly higher for younger than older adults, *t*(60) = 3.27, *p <* .01, *d* = 0.84, and traditional recognition (Figure 2B) did not differ between age groups, *t*(60) = 0.19, *p* = .85, *d* = 0.05. The PDT response probabilities are displayed in Table S1 (bottom rows). Perceptual discrimination (Figure 2C) did not differ between age groups, *t*(60) = 0.12, *p* = .90, *d* = .03. Together, these results replicated the selective age-related mnemonic discrimination deficit consistently shown in object-based MSTs (for a review, see 4) and indicated that perceptual processing abilities could not fully account for this deficit.

**Fig. 2.**
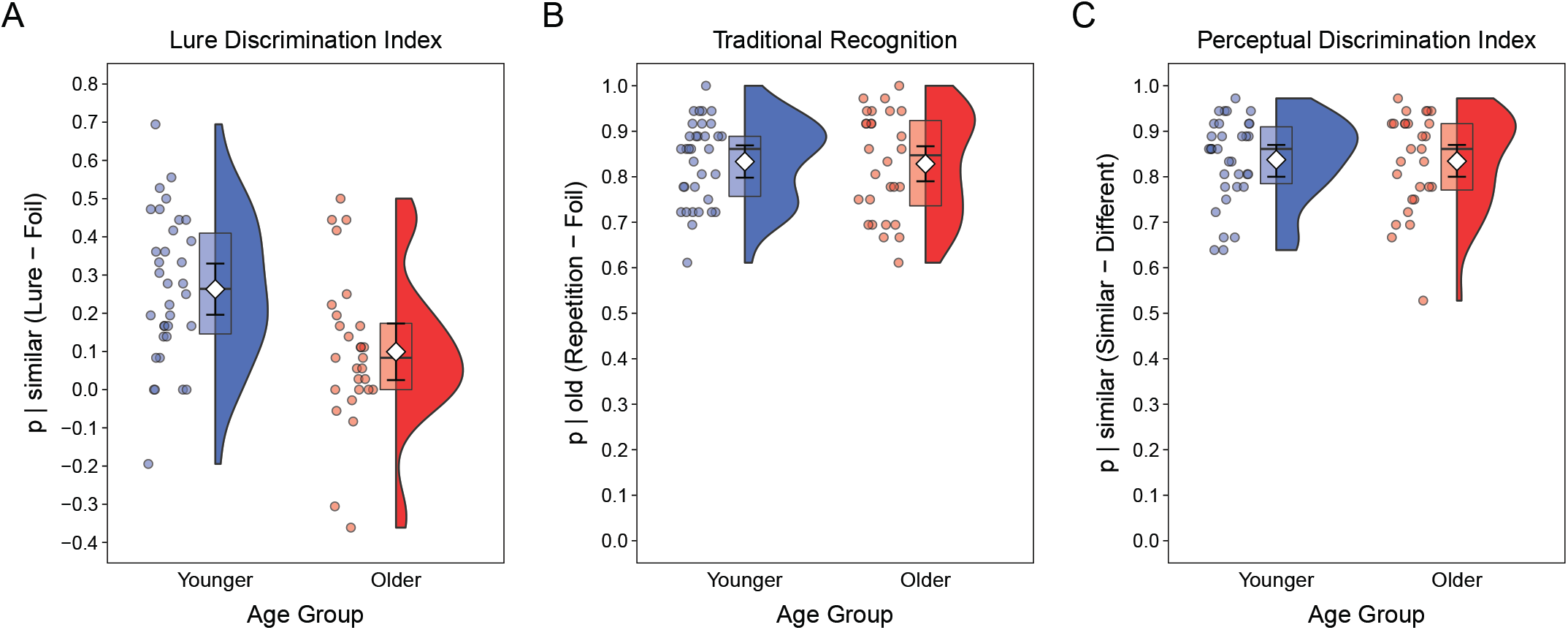
Behavioral Task Performance. (A) Mnemonic discrimination, indicated by Lure Discrimination Index scores, was better for younger than older adults. (B) Recognition, indicated by Traditional Recognition scores, did not differ between age groups. (C) Perceptual discrimination, indicated by Perceptual Discrimination Index scores, did not differ between age groups. (All Panels) Colored points are individual participant probabilities, the widths of the half violin plots represent the proportion of data at each probability, box plots show interquartile ranges and medians, white diamonds are means, and errors bars are 95% confidence intervals.

### Connectome-based Predictive Modeling

#### DMN Connectivity and Mnemonic Discrimination

To test our primary hypothesis that DMN connectivity should predictmnemonic discrimination, we used CPM to correlate predicted with observed LDI scores. Supporting our hypothesis, the positive prediction model was significant, *r*(60) = .39, *p <* .01 (see Figure 3), and 209 out of 903 possible connections (23%) significantly predicted mnemonic discrimination. Controlling for white matter volume alone (instead of total intracranial volume), produced a near-identical predictive relationship, *r*(60) = 0.41, *p <* .01. More generally, these effect sizes are comparable to those reported in previous studies using CPM to predict behavior from intrinsic connectivity (e.g., 70). Because leave-one-subject-out connectome estimations are not independent, a permutation test was performed to assess the likelihood that the positive prediction model emerged by chance. A null distribution was created by randomly shuffling observed LDI scores, re-running CPM, and computing 1000 correlations between predicted and observed scores. There was a significant difference between the null and positive prediction models, *p <* .01, indicating that the positive prediction model was unlikely to have emerged by chance. The negative prediction model, which was not significant, *r*(60) = − 23, *p* = .08, included 22 out of 903 possible significant connections (2%) and is not considered further.

**Fig. 3.**
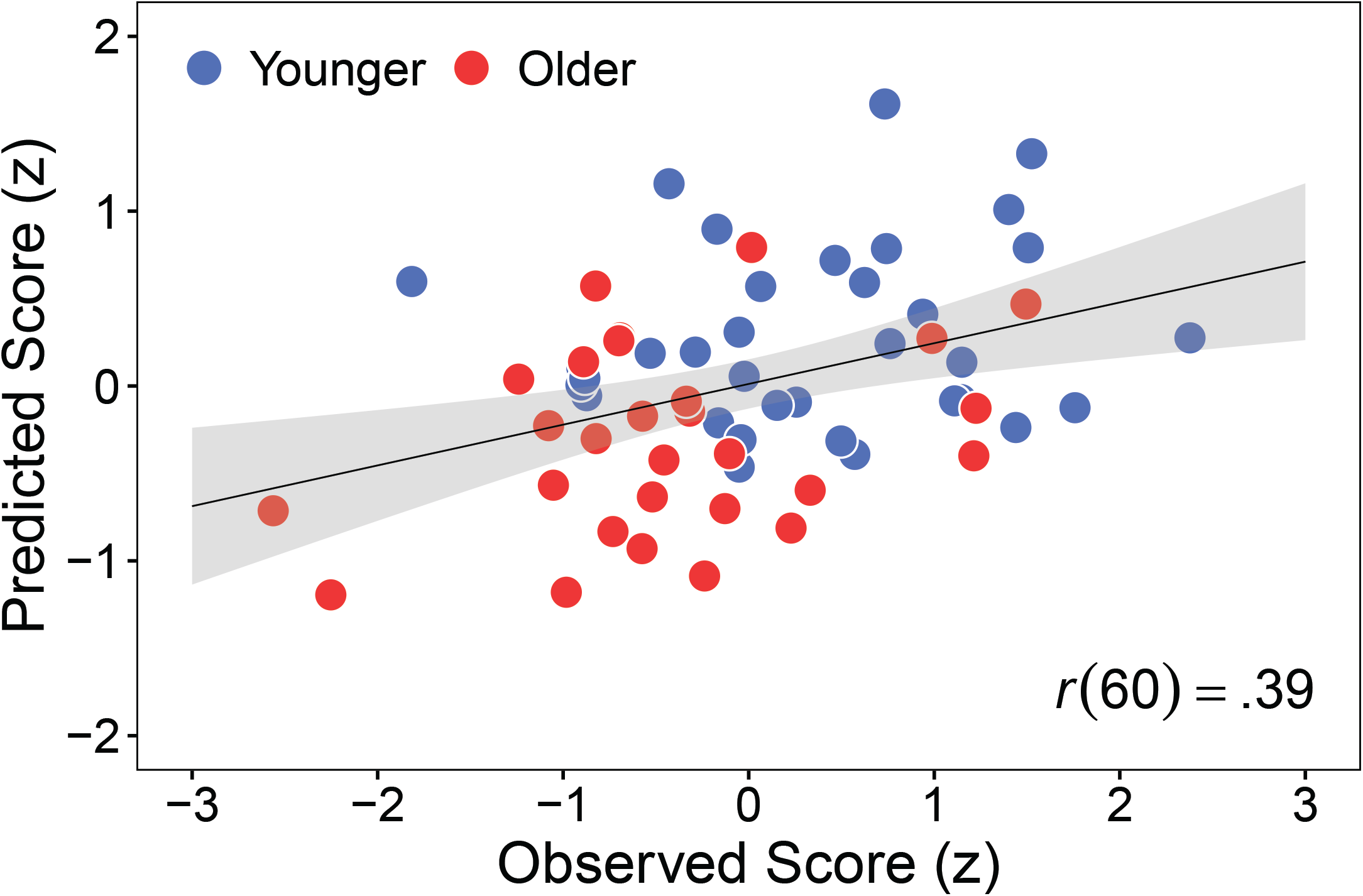
Intrinsic Default Mode Network Connectivity Predicting Mnemonic Discrimination. A scatterplot showing the association between predicted and observed Lure Discrimination Index scores (standardized) based on the Mnemonic Discrimination Connectome for younger (blue) and older (red) adults. Points are individual participants, the line is the best fitting regression line, and the shaded region is the 95% confidence interval.

The connections among regions in the positive prediction model, hereafter referred to as the Mnemonic Discrimination Connectome (MDC), are displayed in Figure 4 (glass brain format) and Figure 5A (grid format). To summarize the locations of these connections, we computed the proportion of connections out of all possible connections within and between four major regions (i.e., prefrontal, hippocampus, temporal, and parietal). In this approach, each proportion is independent of the others. These summaries (Figure 6, top panels) indicate that the relative proportion of connections was greatest within the temporal cortex (panel A) and between the temporal cortex and other regions (panel B). Thus, the temporal regions of the DMN were the most densely connected in the MDC.

**Fig. 4.**
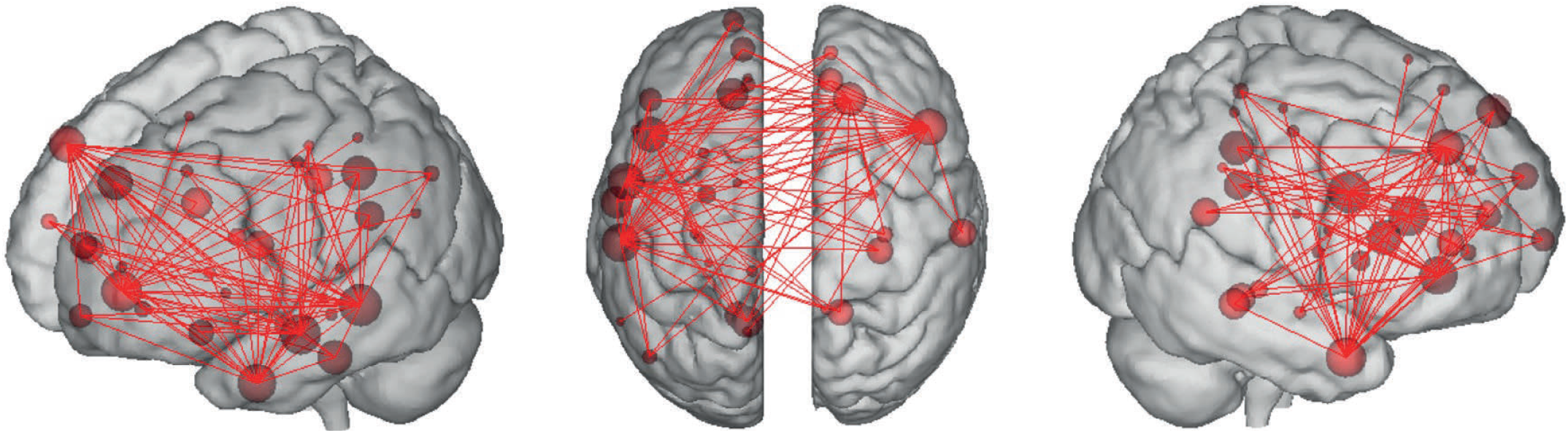
Connections in the Mnemonic Discrimination Connectome (MDC). The MDC visualized from three orientations (left, top, right) using the BioImage Suite Web Viewer: https://bioimagesuiteweb.github.io/webapp/connviewer.html. To provide an interpretable connectome, the threshold for including regions of interest here was set at 20 or more connections. Images using other thresholds can be generated using the data provided on the OSF: https://osf.io/f6vg8/. Larger node sizes (circles) indicate more inter-regional connections (lines).

**Fig. 5.**
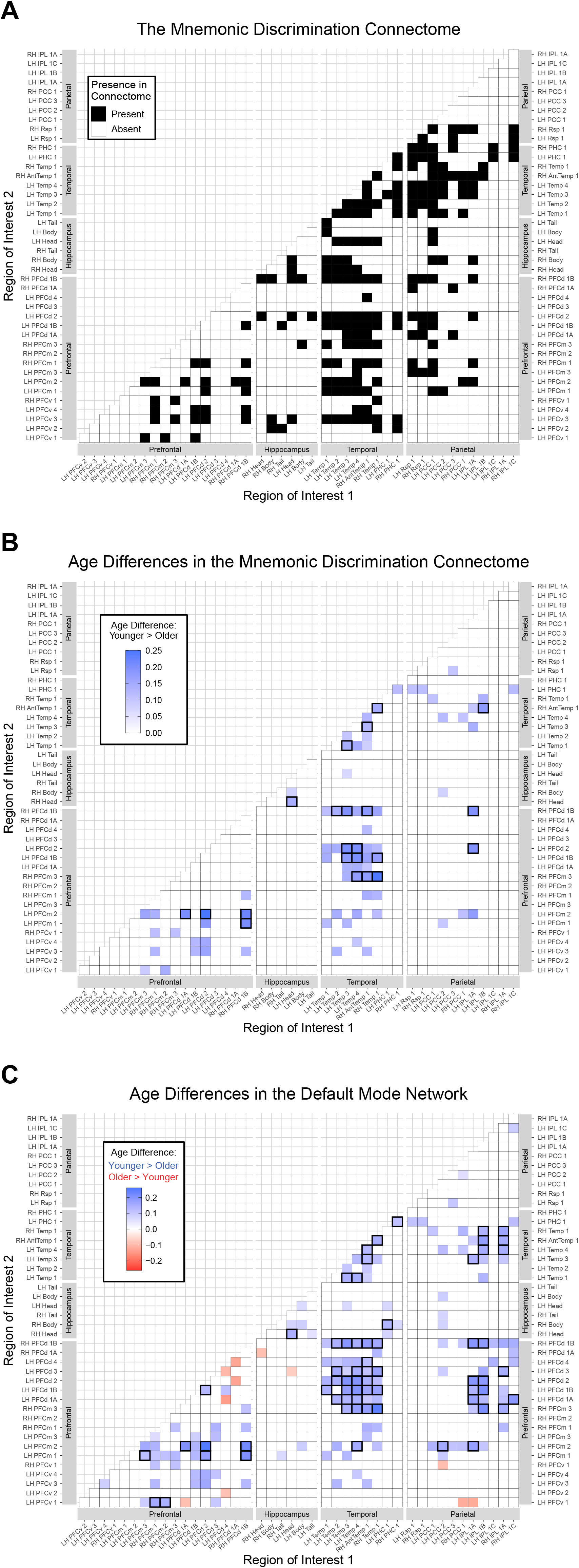
Inter-Regional Connections. (A) The significant connections among regions of interest (ROIs) in the Default Mode Network (DMN) identified in the Mnemonic Discrimination Connectome (MDC). ROI–ROI connections were present in (black squares) or absent from (white squares) the MDC. (B) Age differences in MDC connections. Blue squares show ROI–ROI connections that were stronger for younger than older adults. Connections that were significantly different between age groups and survived False Discovery Rate (FDR) correction are indicated by thicker black borders. (C) Age differences in DMN connections. Blue squares show ROI–ROI connections that were stronger for younger than older adults; red squares show greater connectivity for older than younger adults. Connections that were significantly different between age groups and survived False Discovery Rate (FDR) correction are indicated by thicker black borders. B and C) The color intensities of grid squares indicate the degree of average differences in correlations between Lure Discrimination Index scores and ROI–ROI connections between younger and older adults. (All Panels) ROIs are ordered anterior to posterior starting from the left (x-axis) and bottom (y-axis). Axis labels correspond to the codes in Table 2.

**Fig. 6.**
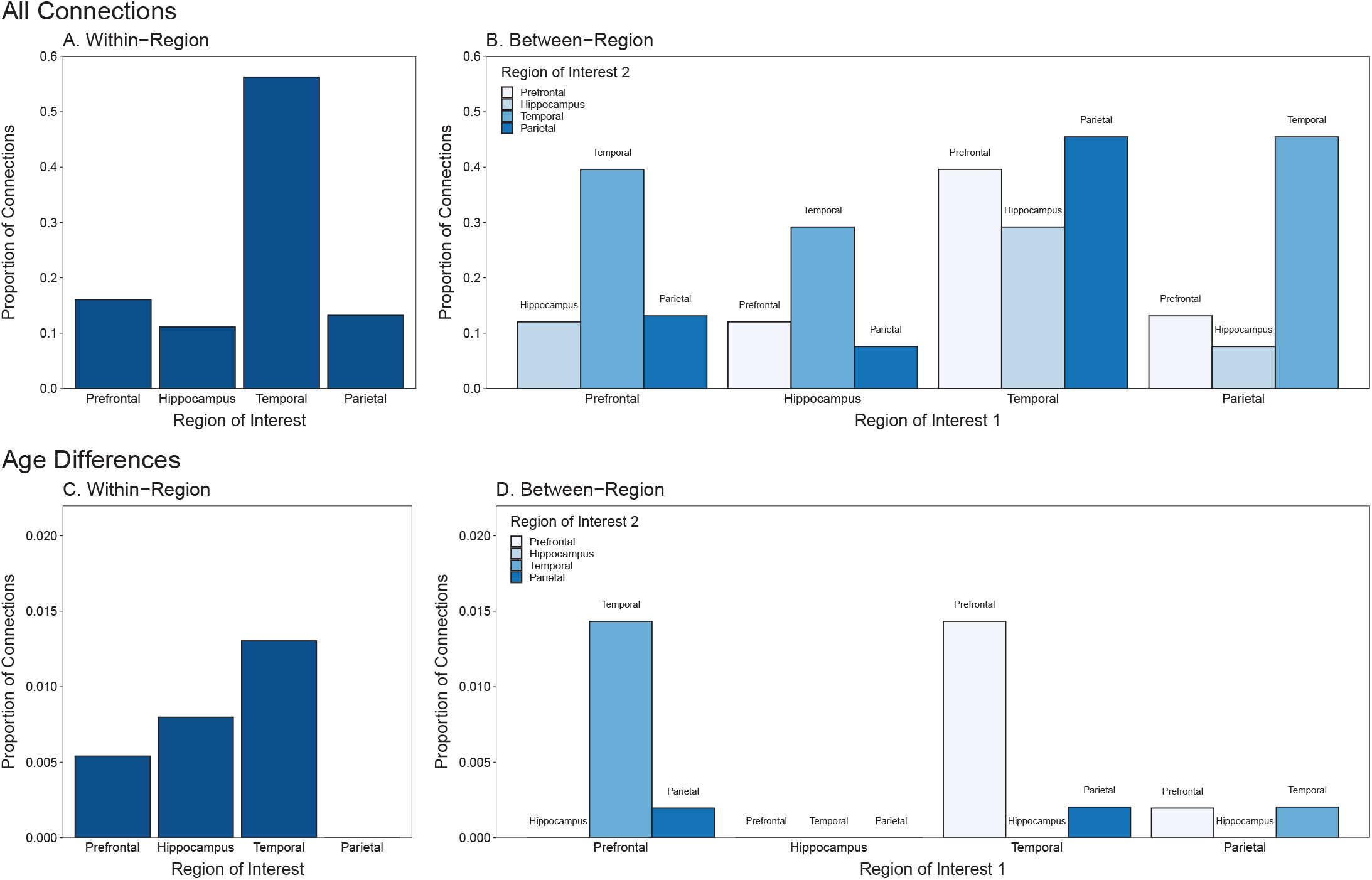
Proportions of Connections in the Major Regions of the Mnemonic Discrimination Connectome. (Top panel) Total proportions of connections within (A) and between (B) regions. (Bottom panel) Proportions of regions showing significantly greater connectivity for younger than older adults that survived false discovery rate correction (C) within and (D) between regions. Proportions were computed separately within each region and between each pair of regions using the total unique number of possible connections in each as the denominator. Therefore, the values above are independent of each other and can all range from 0–1.0.

Next, we established the specificity of intrinsic DMN connectivity in predicting mnemonic discrimination. If the DMN connectivity in the MDC is specifically related to performance on the measure assessing the mechanisms supporting mnemonic discrimination, then the MDC should not predict performance on measures supported by non-identical mechanisms. This outcome would be unsurprising because the mask is defined by its ability to predict mnemonic discrimination (71). This analysis provides an essential sanity check confirming that the MDC was established properly. Figure **ãã** (top panels) shows that the MDC did not significantly predict scores on indices of recognition memory [Panel A, *r*(60) = −0.18, *p* = .15] or perceptual discrimination [Panel B, *r*(60) = 0.11, *p* = .39].

#### Hippocampal Connectivity in the MDC

As described in the Introduction, the MST has primarily been used to examine the relationship between hippocampal function and mnemonic discrimination. To highlight the connectivity involving the hippocampus in the MDC, we explored the interregion connections originating from the head, body, and tail of the hippocampus in both hemispheres (Figure 7). The connectivity was greatest for the left hippocampal head with 11 connections and the right hippocampal body region with 10 connections. Across all divisions of the hippocampus, most connections emerged with temporal (14) and prefrontal (13) regions.

**Fig. 7.**
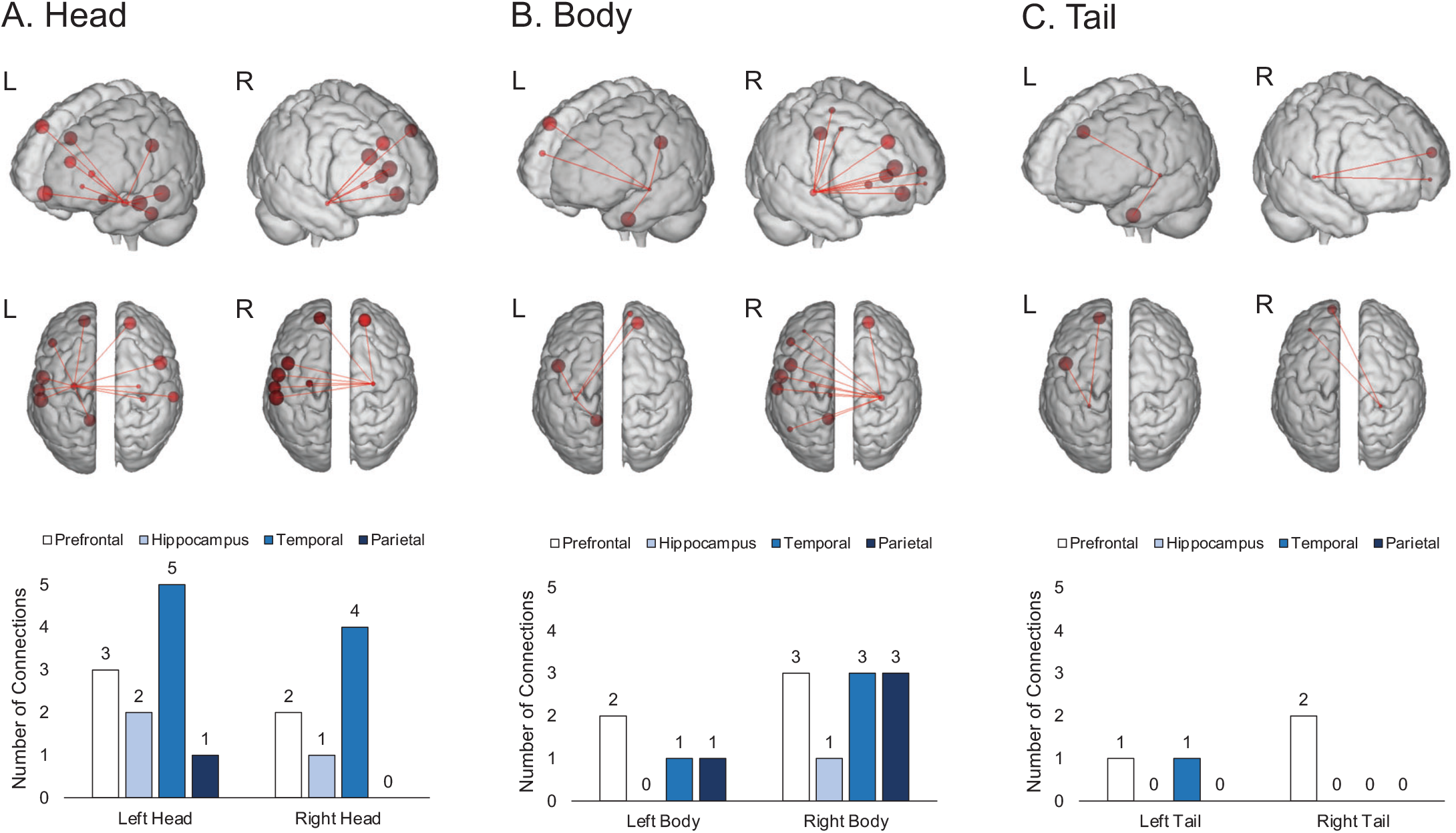
Inter-Region Connectivity from Regions of the Hippocampus in the Mnemonic Discrimination Connectome (MDC). (Top two rows) Connections from the hippocampal head (A), body (B), and tail (C) visualized using the BioImage Suite Web Viewer: https://bioimagesuiteweb.github.io/webapp/connviewer.html. The threshold for including regions of interest here was set at one or more connections. Images using other thresholds can be generated using the data provided on the OSF: https://osf.io/f6vg8/. Larger node sizes (circles) indicate more connections (lines) from each hippocampal region to others in the MDC. (Bottom row) Summaries displaying the numbers of connections from each hippocampal region.

#### Age Differences in MDC Connection Strength

Our secondary aim was to examine if intrinsic DMN connectivity in the CPM predicted mnemonic discrimination more strongly for younger than older adults. We tested for such differences by summing all MDC connections and comparing those values between age groups with a two-sample *t*-test. This measure of connection strength was significantly greater for younger (*M* = 54.05, *SD* = 12.78) than older (*M* = 36.83, *SD* = 14.56) adults, *t*(60) = 4.96, *p <* .001, *d* = 1.26. To determine if this difference was due to chance, we performed a permutation test. We first created a null distribution of *t*statistics based on strength differences between age groups by computing 1000 random connectomes with ROIs including the original MDC connections. The *t*-statistic from the MDC (4.96) was greater than the upper bound of the null distribution (*Range* = 3.64∪4.63, *M* = 4.12, *SD* = 0.19), suggesting that younger adults’ greater connection strength was not likely due to chance.

To identify the locations of age differences in the MDC, we used two-sample t-tests to compare the strength of each ROI–ROI connection between age groups (Figure 5B). Out of the 209 possible connections, 22 (11%) showed greater strength for younger than older adults and survived false discovery rate correction (cells with thicker borders). The stronger connections for younger adults were mostly within the temporal cortex (Figure 6C) and between the temporal and prefrontal cortex (Figure 6D).

Finally, baseline connectivity in the DMN is sometimes stronger for younger than older adults (e.g., 27), suggesting that the age differences in MDC connectivity strength here partly reflected differences in baseline DMN connectivity. We examined the role of DMN connectivity strength in agerelated MDC connectivity differences by comparing ROIROI connection strength in the DMN between age groups (Figure 5C). Younger adults showed significantly stronger average connectivity strength that survived false discovery rate correction in 62 connections between regions across the DMN (cells with thicker borders). Of these differences, 22 (35.5%) were connections within the MDC (Figure 5B). These results suggest that younger adults’ stronger baseline DMN connectivity partly contributed to their stronger connectivity among regions that predicted mnemonic discrimination.

## Discussion

Discriminating existing memories from similar perceptual inputs to mitigate interference is a core feature of episodic memory. Although studies examining the neural mechanisms of mnemonic discrimination have focused primarily on hippocampal structure and function (for a review, see 4), mounting evidence suggests that regions beyond the hippocampus also support this ability (10, 13, 15, 18). The present study extends this nascent literature by characterizing the relationship between functional connectivity among DMN regions and mnemonic discrimination ability. Using a datadriven connectomics approach (34), we showed that interindividual differences in intrinsic connectivity among regions in the DMN predicted mnemonic discrimination. This relationship primarily involved connections with temporal cortex. Further, the better discrimination shown by younger than older adults were primarily reflected in areas where younger adults had stronger temporal-prefrontal connectivity than older adults. These results provide evidence that mnemonic discrimination is supported by connectivity across DMN regions including, but not limited to, the hippocampus. The predictive relationship between DMN connectivity and mnemonic discrimination is consistent with results suggesting that such connectivity supports episodic memory functions (27, 28). DMN subregions, such as medial prefrontal and temporopolar cortex, are presumably involved in mnemonic functions (20, 21). Consistent with these findings, the current study showed that connectivity between such regions predicted mnemonic discrimination. In fact, connections positively related to mnemonic discrimination were broadly distributed across prefrontal, hippocampal, temporal, and parietal regions. These findings suggest that broad cortical connections support mnemonic functions that have often been attributed to hippocampal and adjacent connections in the medial temporal lobes (for a review, see 4). Most of the predictive connections within and between these regions emerged within temporal cortex between regions around the temporal pole. The observed associations between anterior temporal connections and mnemonic discrimination is reminiscent of findings showing that anterior temporal activity is involved in storing semantic representations (72). Relatedly, people with atrophy near and around the temporal pole show difficulty naming and recognizing everyday objects (73, 74). The present findings also suggest that the strength of intrinsic DMN connections involving regions of the anterior temporal cortex may support the ability to distinguish between everyday objects with shared features. These findings accord with views emphasizing that anterior temporal regions mediate the representations of object identities (23), and extends such views by emphasizing the importance of functional interactions among regions that do not selectively support a single task. Importantly, we do not interpret the present findings as suggesting that pattern separation computations occur in extrahippocampal cortical regions per se. Instead, we suggest that networks comprising cortical regions that communicate with the hippocampus may indirectly support pattern separation performed by hippocampal regions. This could occur, for example, if such cortical-hippocampal networks support the encoding of high-fidelity representations that enable comparison with and rejection of similar lures via a recall-to-reject mechanism (cf. 1, 11).

Following this suggestion, one could predict a key role for connectivity between the hippocampus and parietal regions in mnemonic discrimination. Indeed, the prominent structural and functional connectivity between the hippocampus and parietal regions has led some to suggest that there is a ‘parietal memory network’ within the DMN that supports episodic memory (75, 76). Although network connections of this kind could potentially enable a recall-to-reject mechanism, the present results showed greater connectivity strength of predictive connections within more anterior than posterior DMN regions. These findings are consistent with accounts positing that an anterior-temporal DMN subnetwork preferentially supports memory for items and features of items (19, 23, 31, 77), and implies that memory for contextual features associated with items representations supported by posterior DMN connections (cf. 23) may make lesser contributions to processes leading to mnemonic discrimination, at least in task variants using objects as stimuli.

Consistent with the view that hippocampal pattern separation processes may interact with cortical processes to support mnemonic discrimination, weaker functional connectivity between the anterior hippocampus and parahippocampal cortex during MST performance has been associated with poorer mnemonic discrimination in older adults (33). The present findings are consistent with this result in that functional connectivity between the anterior hippocampus and cortical regions of the DMN showed the clearest age-related differences. However, differing from earlier findings, the present results suggest that intrinsic relationships in activity among DMN regions, even at rest, predict the ability to discriminate sensory inputs from stored representations in the service of preventing catastrophic memory interference. The prominent anterior cortical connections in the MDC suggest that functional interactions between anterior temporal and prefrontal regions may play critical roles in mnemonic discrimination ability and have important interactions with anterior portions of the hippocampus. The nature and extent of these relationships are ideal targets for future studies of mnemonic discrimination.

It is also noteworthy that DMN connections predicted mnemonic discrimination but did not predict perceptual discrimination or traditional recognition. The absence of DMN connections predicting perceptual discrimination suggests more specific DMN involvement in mnemonic processes than visual processing of feature differences between objects. Given that traditional recognition is clearly an aspect of episodic memory, it is perhaps surprising that DMN connections did not predict that ability. The processes involved in mnemonic discrimination and traditional recognition are not identical, however, suggesting that specific types of episodic memory processes are supported by intrinsic DMN connectivity. Indeed, the exact neural mechanisms underlying these memory processes are likely distinct (78). One possibility to be explored in future research is whether intrinsic connections between the DMN and other brain networks predict traditional recognition. Consistent with this possibility, intranetwork connections have been linked to cognitive abilities, such as verbal and visual creativity (79).

The present study provides an innovative contribution to the mnemonic discrimination literature because it is the first to our knowledge to use a connectomics approach to identify the neural mechanisms associated with individual differences in mnemonic discrimination ability. The CPM approach enabled exploratory analyses extending beyond traditional whole-brain activation patterns. Although CPM is usually applied to whole-brain connectivity (34), we took a more targeted, theoretically motivated approach by examining functional connectivity among DMN regions including the hippocampus. Importantly, this reduced the number of ROIs in the connectome, thus allowing for more precise assessments of brain-behavior associations, such as identifying age differences in mnemonic discrimination ability predicted by connectivity strength in anterior, medial, and posterior regions of the DMN.

The greater predictive strength of the MDC for younger than older adults suggested that older adults have less defined DMN co-activation predicting mnemonic discrimination. Older adults also showed weaker overall baseline DMN connectivity than younger adults, including the connections in the MDC. Dysfunctional circuitry among DMN regions associated with healthy aging is also associated with episodic memory deficits (3, 30–32). The present findings thus suggest that poorer baseline network cohesion may be less predictive of behavior under the CPM approach.

The above-described MDC revealed DMN connections predictive of the wide range of mnemonic discrimination performance expected across younger and older adults. Similar methods have been recently used to capture connectomes predicting behavior across different groups (e.g., 39). Other recent studies have included group-distinguishing covariates in CPM to determine connections predictive of behavior beyond group differences (e.g., 39, 80). Our approach combined these methods. Whereas the above-described MDC revealed overall predictive connections, an age-controlled MDC revealed connections predicting mnemonic discrimination that should not be attributable to age. The fact that an age-controlled CPM significantly predicted mnemonic discrimination indicates that age was not the only reason why the original MDC emerged.

We also found that the age-controlled MDC connections that overlapped with the original MDC connections comprised over half the total original connections. Thus, the majority of the predictive connections in the MDC could not be solely attributed to age differences. In addition, almost 90% of the connections in the age-controlled MDC were present in the original MDC. This indicated that the age-controlled MDC was largely a subset of the original MDC and not a completely independent set of predictive DMN connections. Future work could compare original to group-controlled MDCs to further identify which predictive connections are driven primarily by group status. However, there is not yet a standardized approach for making such comparisons (see e.g., 39).

A key translational application of the CPM approach is to detect cognitive impairments associated with Alzheimer’s disease from patterns of intrinsic whole-brain connectivity (81). The MDC established here in a visual object recognition paradigm could serve as a biomarker of normal and pathological cognitive decline. This possibility is based on research showing that impaired visual object discrimination is associated with mild cognitive impairment and Alzheimer’s disease (82) as well as beta-amyloid protein deposition, which is an earlier indicator of Alzheimer’s disease pathology (83). If the MDC predicts Alzheimer’s risk in older adults, it could potentially be used as an early indicator of eventual conversion. This direction is especially promising if neural predictors of mnemonic discrimination ability can be obtained from brief resting-state scans.

The present study had some limitations. First, resting-state functional connectivity data is susceptible to contamination from uncontrolled sources of variability (e.g., participant respiration). Although we accounted for this using a conservative preprocessing method that partly controls for such variability (59), future work could also include physiological measures as regressors in comparable analyses. Second, although the current sample size is comparable to related work (16), more reliable estimates between intrinsic connectivity and behavior can be derived from larger samples than we tested here (84, 85). Finally, although our focus on DMN connectivity was theoretically motivated, communication among networks may also predict mnemonic discrimination (e.g., (38)). Given that mnemonic discrimination requires visual acuity, future CPM approaches could include multiple networks (e.g., sensory and attention networks with the DMN) to identify between-network interactions that predict mnemonic discrimination ability.

To conclude, the present study was the first to leverage a connectomics approach to examine the association between individual variation in intrinsic functional connectivity among DMN regions and mnemonic discrimination of visual objects. The MDC established here showed that intrinsic connections within and between temporal, prefrontal, parietal, and hippocampal regions at rest predicted mnemonic discrimination ability. Such relationships indicate clearly that a comprehensive explanation of the neural mechanisms of mnemonic discrimination requires widening the focus beyond the medial temporal lobes and adjacent subcortical structures to examine broader cortical networks including those structures.

## ACKNOWLEDGEMENTS

The data that support the findings of this study are openly available in the Open Science Framework at https://osf.io/f6vg8/ and OpenNeuro at https://doi.org/10.18112/openneuro.ds003871.v1.0.2.

