## Supplementary material for "Intrinsic Functional Connectivity in the Default Mode Network Predicts Mnemonic Discrimination: A Connectome-based Modeling Approach": WCRC_MST_Connectivity_Hipp_SI.pdf

### Supplemental Information

#### 1. Connectome-based Predictive Modeling

As indicated in the main manuscript, we used connectome-based predictive modeling (CPM) to create a mnemonic discrimination connectome (MDC) using ROI-ROI connections in the default mode network (DMN) controlling for intracranial volume and including age as a continuous covariate (described below).

##### *DMN Connectivity and Mnemonic Discrimination*

To test our primary hypothesis that DMN connectivity should predict mnemonic discrimination, we used CPM to correlate predicted with observed lure discrimination index (LDI) scores. The positive prediction model was significant, $r(60) = .45$, $p < .001$ (Figure S1), and 129 out of 903 possible connections (14%) significantly predicted mnemonic discrimination. Controlling for white matter volume alone (instead of total intracranial volume), produced a near-identical predictive relationship, $r(60) = 0.45$, $p < .001$. A permutation assessed the likelihood that the positive prediction model emerged by chance. A null distribution was created by randomly shuffling observed LDI scores, re-running CPM, and computing 1000 correlations between predicted and observed scores. There was a significant difference between the null and positive prediction models, $p < .01$, indicating that the positive prediction model was unlikely to have emerged by chance. The non-significant negative prediction model, $r(60) = -.22$, $p = .09$, included 32 out of 903 possible significant connections (4%) and is not considered further.

The connections among regions in the positive prediction model (i.e., the MDC), are displayed in Figure S2 (glass brain format) and Figure S3 (grid format). Of the 129 connections that significantly predicted mnemonic discrimination in this model, which included age as a covariate, 116 (89.9%) connections were present in the original CPM that did not control for age differences. There were also 13 unique connections in the age-controlled CPM that were not in the original CPM. In addition, the 116 connections in the age-controlled CPM that overlap with the original CPM comprised 55.5% of the 209 connections in the original CPM. To summarize the locations of these connections, we computed the proportion of connections out of all possible connections within and between four major regions (i.e., prefrontal, hippocampus, temporal, and parietal). In this approach, each proportion is independent of the others. These summaries (Figure S4) indicate that the relative proportion of connections was greatest within the temporal cortex (panel A) and between the temporal cortex and other regions (panel B). Thus, although controlling for age resulted in an MDC with fewer overall ROI-ROI connections that significantly predicted mnemonic discrimination than the original MDC, the temporal regions of the DMN were still the most densely connected.

Next, we established the specificity of intrinsic DMN connectivity in predicting mnemonic discrimination. If the DMN connectivity in the MDC is specifically related to performance on the measure assessing the mechanisms supporting mnemonic discrimination, then the MDC should *not* predict performance on measures supported by non-identical mechanisms. This outcome would be unsurprising because the mask is defined by its ability to predict mnemonic discrimination (Kriegeskorte et al., 2009). This analysis provides an essential sanity check confirming that the MDC was established properly. Figure S5 (top panels) shows that the MDC did not significantly predict scores on indices of recognition memory [Panel A, $r(60) = -.14$, $p = .27$] or perceptual discrimination [Panel B, $r(60) = .17$, $p = .17$].

We also examined whether intrinsic DMN connectivity selectively predicted mnemonic discrimination by applying CPM to recognition and perceptual discrimination scores. This established whether a behavior-specific could be identified for such indices. Given our primary focus on the significant positive prediction model for LDI scores, we focus here only on positive prediction models for recognition and perceptual discrimination. Figure S5 (bottom panels) shows that connectomes built from intrinsic DMN connectivity did not have sufficient explanatory power to predict scores on recognition [Panel C, $r(60) = .01$, $p = .91$] or perceptual discrimination [Panel D, $r(60) = .14$, $p = .28$] indices. Collectively, these results indicate that intrinsic DMN connectivity selectively predicted mnemonic discrimination even when controlling for age differences.

**Table S1**

*Response Probabilities on the Mnemonic Similarity and Perceptual Discrimination Tasks*

| | Mnemonic Similarity Task (Test Objects) | | | | | |
|---|---|---|---|---|---|---|
| | Similar Lures | | Repeated Objects | | Novel Foils | |
| Response | Younger | Older | Younger | Older | Younger | Older |
| Similar | **.38 [.34, .42]** | **.23 [.18, .27]** | .08 [.06, .10] | .07 [.04, .09] | .12 [.08, .15] | .13 [.09, .16] |
| Old | .53 [.50, .57] | .62 [.59, .67] | **.87 [.85, .89]** | **.86 [.84, .88]** | .04 [.01, .07] | .03 [.01, .07] |
| New | .07 [.03, .10] | .08 [.04, .12] | .03 [.01, .05] | .04 [.02, .06] | **.82 [.79, .86]** | **.75 [.72, .79]** |
| Missing | .02 [.00, .06] | .07 [.03, .11] | .02 [.00, .04] | .03 [.01, .06] | .02 [.00, .05] | .09 [.05, .12] |

| | Perceptual Discrimination Task (Object Pairs) | | | | | |
|---|---|---|---|---|---|---|
| | Similar Objects | | Repeated Objects | | Novel Objects | |
| Response | Younger | Older | Younger | Older | Younger | Older |
| Similar | **.88 [.86, .91]** | **.85 [.82, .88]** | .07 [.05, .09] | .06 [.05, .09] | .05 [.03, .07] | .02 [.00, .04] |
| Same | .08 [.05, .10] | .09 [.06, .12] | **.93 [.91, .94]** | **.93 [.91, .95]** | .01 [.00, .03] | .01 [.00, .03] |
| Different | .04 [.01, .06] | .06 [.03, .09] | .00 [.00, .00] | .01 [.00, .03] | **.94 [.92, .96]** | **.97 [.95, .99]** |

*Note.* Bold values are correct response probabilities. Brackets contain 95% confidence intervals.

### Figure Captions

**Figure S1.** Intrinsic Default Mode Network Connectivity Predicting Mnemonic Discrimination Including Age as a Covariate**.** A scatterplot showing the association between predicted and observed Lure Discrimination Index scores (standardized) based on the Mnemonic Discrimination Connectome for younger (blue) and older (red) adults including age as a covariate in the model. Points are individual participants, the line is the best fitting regression line, and the shaded region is the 95% confidence interval.

**Figure S2.** Connections in the Mnemonic Discrimination Connectome (MDC) Including Age as a Covariate. The MDC visualized from three orientations (left, top, right) using the BioImage Suite Web Viewer: https://bioimagesuiteweb.github.io/webapp/connviewer.html. To provide an interpretable connectome, the threshold for including regions of interest here was set at 15 or more connections. Images using other thresholds can be generated using the data provided on the OSF: https://osf.io/f6vg8/. Larger node sizes (circles) indicate more inter-regional connections (lines).

**Figure S3.** Inter-Regional Connections for the Age-Controlled and Original Mnemonic Discrimination Connectomes (MDCs). (A) The significant connections among regions of interest (ROIs) in the Default Mode Network (DMN) identified in two MDCs. The significant connections in the age-controlled MDC appear as black squares for connections present in both the age-controlled and original MDCs and as dark gray squares for connections present only in the age-controlled MDC. Connections from the original MDC that were not present in the age-controlled MDC appear as light gray squares. Connections that were absent from both MDCs appear as white squares. ROIs are ordered anterior to posterior starting from the left (x-axis) and bottom (y-axis). Axis labels correspond to the codes in Table 2.

**Figure S4.** Proportions of Connections in the Major Regions of the Mnemonic Discrimination Connectome Including Age as a Covariate. Total proportions of connections within (A) and between (B) regions. Proportions were computed separately within each region and between each pair of regions using the total unique number of possible connections in each as the denominator. Therefore, the values above are independent of each other and can all range from 0-1.0.

**Figure S5.** Intrinsic Functional Default Mode Network Connectivity Predicting Recognition and Perceptual Discrimination Including Age as a Covariate. Scatterplots showing the association between predicted and observed Traditional Recognition (A and C) and Perceptual Discrimination Index (B and D) scores (standardized). (Top panel) Associations including predicted scores based on the Mnemonic Discrimination Connectome. (Bottom panel) Associations including predicted scores based on connectomes created for recognition and perceptual discrimination based on intrinsic default mode network connectivity. Points are individual participants (younger = blue; older = red), the lines are the best fitting regression lines, and the shaded regions are 95% confidence intervals.
